# Oscillations synchronize amygdala-to-prefrontal primate circuits during aversive learning

**DOI:** 10.1101/184531

**Authors:** Aryeh Taub, Rony Paz

## Abstract

The contribution of oscillatory synchrony in the primate amygdala-prefrontal pathway to aversive learning remains unknown. We found increased power and phase synchrony in the theta range during aversive conditioning. The synchrony was linked to single-unit spiking and exhibited specific directionality between input and output measures in each region. Although it was correlated with the development of conditioned responses, it declined once the association stabilized. The results suggest that amygdala spikes aid to synchronize ACC activity and transfer error-signal information to support memory formation.

**Highlights:**

- Tone-odor conditioning induces theta phase-reset in primate amygdala and dACC
- A directional phase-locking develops between amygdala spikes and dACC Theta
- Information transfer from Amygdala to dACC decreases once memory stabilizes

## Introduction

Neuronal oscillations are considered a mechanism that synchronizes spiking activity across brain regions and can enhance information transfer (Buzsáki et al., 2012; Deffains et al., 2016). In rodents, much evidence suggests that oscillations at the Theta range (4 – 8 Hz) support amygdala-prefrontal interactions and contribute to expression (Courtin et al., 2014), discrimination (Likhtik et al., 2014) and consolidation (Popa et al., 2010) of fear memories. In primates, spiking synchrony underlies persistent aversive memories (Livneh and Paz, 2012), and successful updating between fear and safety (Klavir et al., 2013). Yet it remains unclear if neural oscillations, and specifically at the theta range, contribute to spiking synchrony in the primate amygdala-prefrontal pathway and in turn to successful formation of aversive memories. Moreover, it is unknown what is the dynamics of such mechanism during learning, and what kind of information this synchrony carries.

We therefore examined the emergence of theta and resetting of phase in the amygdala and anterior-cingulate-cortex (dACC) of monkeys while they acquire novel aversive tone-odor associations on a daily basis. We further tested the relation between theta phase synchrony and single-unit activity in both regions, and the directionality of information transfer during learning between the two regions. We provide here first evidence for the role of theta in the primate amygdala-PFC circuitry and its dynamical role during aversive learning.

## Results

### Increased theta power in the primate amygdala and dACC

We examined the development of theta oscillations in the amygdala and dACC of monkeys while they acquire novel aversive tone-odor associations on a daily basis (n_days_ = 19 + 13, Fig. 1a-c). We found increased power following the CS and before delivery of the aversive unconditioned stimulus (US) in both regions (Fig. 1d-e, t-test for mean power in theta during 1 sec following CS onset vs. mean of 5 consecutive seconds during the pre-CS, *n*_amygdala_ = 73, *p*_amygdala_ = 0.007, Δ(CS/Baseline_amygdala_) = 1.18, *n*_dACC_ = 76, *p*_dACC_ = 1.6^−13^, Δ(CS/Baseline_dAee_) = 1.66). In keeping with this, autocorrelation of single-units revealed an increased component between 4 - 8 Hz (t-test, *n*_amygdala_ = 214, *P*_amygdala_ = 2^−9^, Δ(CS/Baseline_amygdala_) = 2.85, *n*_dACC_ = 217, *P*_dACC_ = 5^−6^, Δ(CS/Baseline_dAee_) = 2.54, Fig.1f). Interestingly, the power increased throughout conditioning (Fig. 2a and inset, one way ANOVA, *F_amygdala_* = 4.54, *df_amygdala_* = 4, *P*_amygdala_ = 0.005, *F*_dACC_ = 4.8, *df*_dACC_ = 4, *p_dACC_* = 0.007) with increased theta power between habituation and conditioning trials 11 - 20 (post-hoc t-test, *p* < 0.001). Taken together, increased power of theta oscillations and pattern of spiking discharge suggest a role for theta activity in the primate amygdala and dACC.

**Figure 1.**
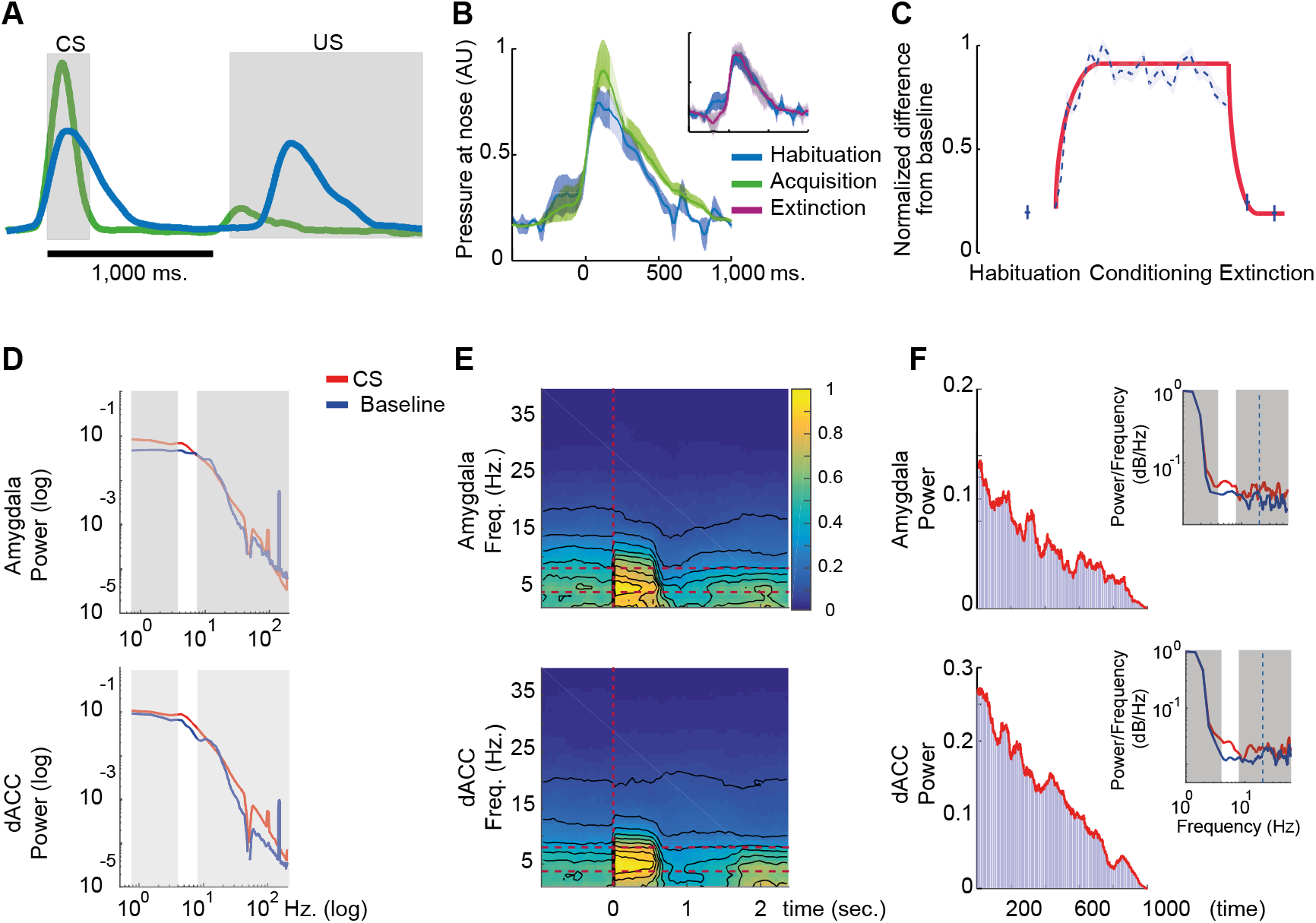
Increased theta power in response to auditory stimulus. (A) Paired conditioning trials (green line): real-time detection of inhale elicited the presentation of a conditioned stimulus (CS, a pure tone) that predicted the release of unconditioned stimulus at the onset of the next inhale (US, an aversive odor, left and right gray rectangles mark the durations of CS and US, respectively). Aversive odors resulted in decreased inhale (UR), and increased inhale to the tone (CR). During habituation and extinction trials (blue line), the CS was not followed by a US. Shown is real data taken from one trial. (B) Following conditioning trials (green line), mean size of the CR increased compared to both habituation (blue) and extinction (maroon), with the latter two not significantly different from each other. (C) Increased CRs, measured as area under the curve during 350 ms post-CS compared with mean inhale volume during pre-CS time (i.e. taken during the ITI) was observed during acquisition trials, with return to baseline levels during extinction trials. The red line is a fit for presentation purposes only. (D) An example of power spectral density analysis indicating an increase in theta power in response to the CS. Data is presented as log/log power with white band (nonshaded) highlighting the theta band. (E) Theta power increases in response to the CS, mean over all sessions and electrodes (horizontal doted red lines mark theta band range). (F) Example of autocorrelation of single-units recorded from the amygdala (top) and dACC (bottom) showing an increase in power between 4 - 8 Hz (insets in log/log, Theta band is presented with white non-shaded background for during CS (solid red) and during intertrial interval baseline activity (solid blue). Blue dashed represents line indicates for 20 Hz.

**Figure 2.**
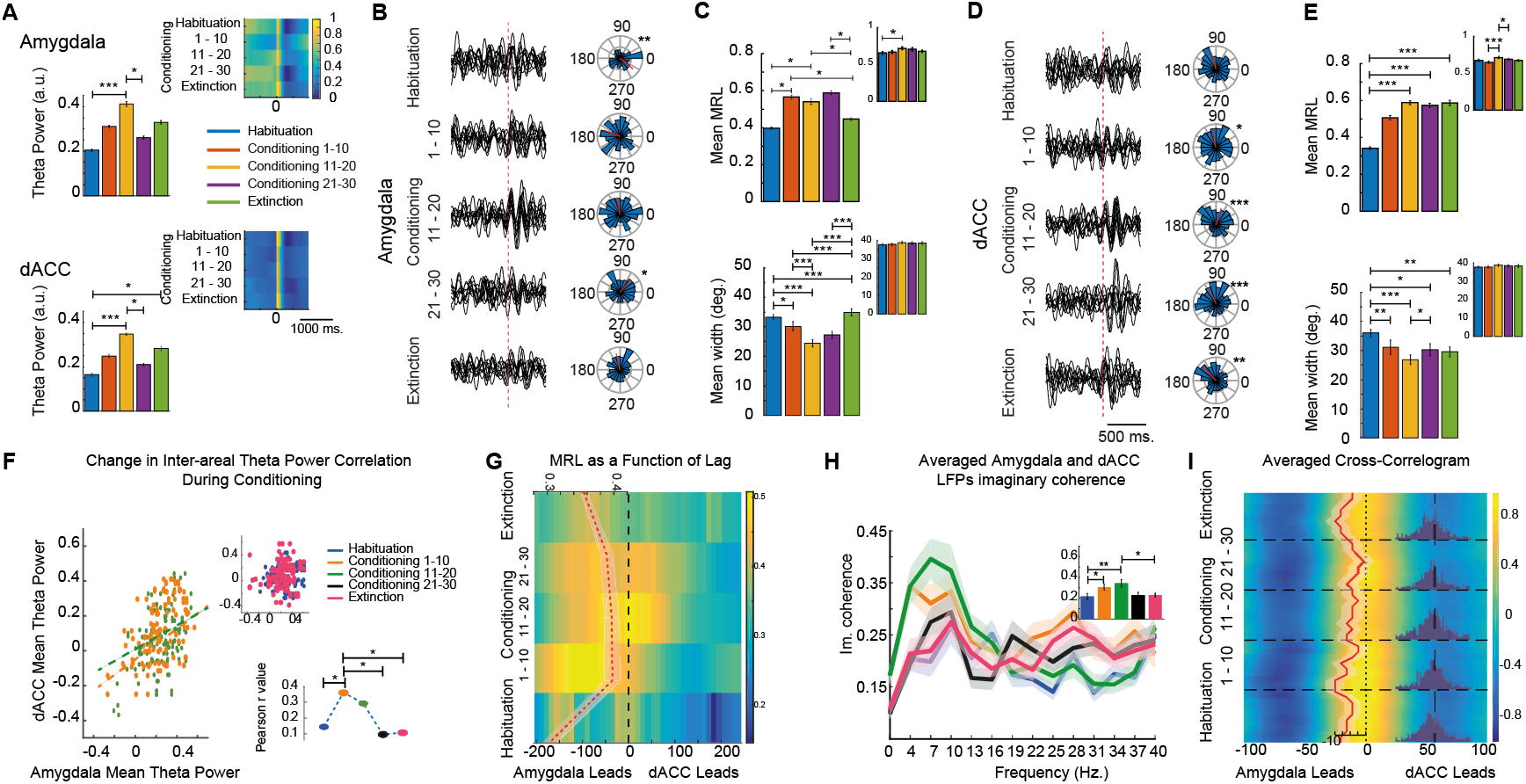
Conditioning induces increased theta power and phase resetting. (A) Mean theta power during 1000 ms following CS onset increased significantly during conditioning. Inset illustrates mean theta power during 1000 before and following CS onset along a daily session of habituation, conditioning and extinction. (B) Phase reset developed during conditioning in both the amygdala and dACC (D). Insets presents the distribution and mean (red line) of theta phases across electrodes and days of recordings. (C) Phase reset was characterized by increased MRL score and decreased half-width theta-phase distribution in the amygdala and dACC (E), with some change in MRL score but not half-width phase distribution during pre-CS period (insets). (F) Within-trial correlation in theta power between the amygdala and dACC increases during initial stages of conditioning, and then decreases during the last ten trials of conditioning (not different than habituation and extinction, top inset). Bottom inset shows the change in correlation values along conditioning. (G) Phase synchrony as a function of lag between the two structures. Red line indicates the mean MRL score ± SEM and time of maximal MRL. (H) Imaginary coherence reveals specificity of conditioning dependent increase in LFP power to the theta band, and to trials 1-21. (I) Mean cross-correlation from 1000ms post-CS (red line indicates the mean, Signed rank test of difference from zero, *p* < 0.0001). Insets show the histogram of crosscorrelation peaks. The amygdala leads (−7.7 ms, p < 1^−10^, signed rank test). * *p* < 0.05, ** *p* < 0.01, *** *p* < 0.001.

### Tone – odor conditioning induces phase resetting at the Theta range

It was recently suggested that theta phase-resetting can be used to organize local spiking activity and drive fear expression in mice (Courtin et al., 2014; Karalis et al., 2016; Likhtik et al., 2014). In the primate, we observed significant phase resetting which developed in response to the tone during conditioning (Fig. 2b-e-f). We found increased synchronization and decreased half-width theta-phase distribution in the amygdala *(n* = 42, one way ANOVA, *F_Synchronization_* = 16.5, *df_Synchronization_* = 4, *P*synchronization = 3^−11^, *F_haif-width_* = 19.18, *df_haif-width_* = 4, *p_haif-width_* = 2^−13^) as well as in the dACC (*n* = 57, *F_Synchronization_* = 22.7, *df_Synchronization_* = 4, *P_synchronization_* = 2^−15^, *F_haif-width_* = 13.92, *df_haif-width_* = 4, *P_haif-width_* = 2^−10^), with specific changes between habituation and conditioning trials 11 - 20 (post-hoc t-test, *P*_Synchronization_ < 0.03 and *p_haif-width_* < 0.001, Bonferroni corrected). In comparison, the same measure was not significant during the intertrial interval (*p_ITI amygdaia_* > 0.5, *P_iTi dACC_* > 0.5).

### Conditioning increases amygdala-dACC theta synchorny

Next, to examine functional connectivity between the amygdala and the dACC during conditioning trials, we computed and found an increase in the correlation of theta power across the two structures (Fig. 2f; Spearman’s correlation, *n* = 129, *r_habituation_* = 0.14, *P_habituation_* = 0.1, *r_conditioning 1-10_* = 0.36, *P_conditioning 1-10_* = 0.0001), with increase from habituation to conditioning trials 1-10 and 11-20 (Fig. 2f insets, *p* < 0.05 for both). The increased functional connectivity hypothesis was further supported by increased phase synchronization between the amygdala and dACC. Specifically, synchronization was calculated as a function of the lag (Fig. 2g, two way ANOVA, for *F*_conditioning stage_ = 14.22, *df*_conditioning stage_ = 4, *P*_conditioning stage_ = 3^−9^, *F_iag_* = 5.02, *df_iag_* = 40, *P_iag_* = 8^−14^), with interaction of stage by lag (*F* = 1.37, *df* = 160, *p* = 0.001), but not during pre-CS activity (two way ANOVA, for *F*_conditioning stage_ = 0.96, *df*_conditioning stage_ = 4, *P*_conditioning stage_ = 0.43). To examine the band specificity of the phase synchronization, we used imaginary coherence analysis and found a specific increase in theta coherence during conditioning (Fig. 2h, two way anova, *F_band_* = 7.48, *df_band_* = 21, *p_band_* = 0.001, *F*_stage_ = 7.48, *df_stage_* = 3, *P_stage_* = 0.0001), with band by stage interaction effect (*F* = 1.41, *df* = 63, *p* = 0.02), and specific increase during conditioning trials 1 -10 and 11 – 21 (post-hoc, *p* < 0.05). No differences were observed in imaginary coherence during pre-CS period (two way anova, *F_band_* = 1.06, *df_band_* = 21, *p_band_* = 0.39). Taken together, the results suggest that theta oscillations can be used as long-range communication between the amygdala and the dACC during conditioning in the primate. Finally, the lag and negative peak of cross-correlation (Fig. 2i) suggests a directional flow of information from the amygdala to the dACC.

### Phase locking develops between amygdala-spikes and dACC-theta

To further examine the directionality of amygdala-dACC interactions, we quantified the synchrony between the output of one region - measured as spiking activity, and the input in the other region - approximated by the theta oscillations (Buzsáki et al., 2012) (Fig. 3a-e). This revealed a phase-locking between amygdala spikes and dACC oscillation, but not vice versa (Fig. 3b). Specifically, the proportion of amygdala units that were locked to theta oscillations in the dACC increased during conditioning by about 35% (n = 234, from 11.25% to 17.4%), while there was no increase in synchronization during the intertrial interval and for the shuffled data (both < 0.6% significant cells throughout conditioning stages). In these pairs of electrodes, synchronization score significantly increased during conditioning (3ig. 4c, one way anova, *F* = 4.34, *df* = 4, *p* = 0.002), with increased synchronization during trials 11 - 20 and 21 - 30 (post-hoc *p* = 0.002 *and p* = 0.003, respectively). Spike triggered LFP analysis revealed same results (Fig. 3d). However, the proportion of dACC units that were locked to theta oscillations in the amygdala decreased by about 32% (Fig. 3b inset, *n* = 289, from 12.32% to 8.33%) and synchronization score did not increase significantly compared to habituation levels.

**Figure 3.**
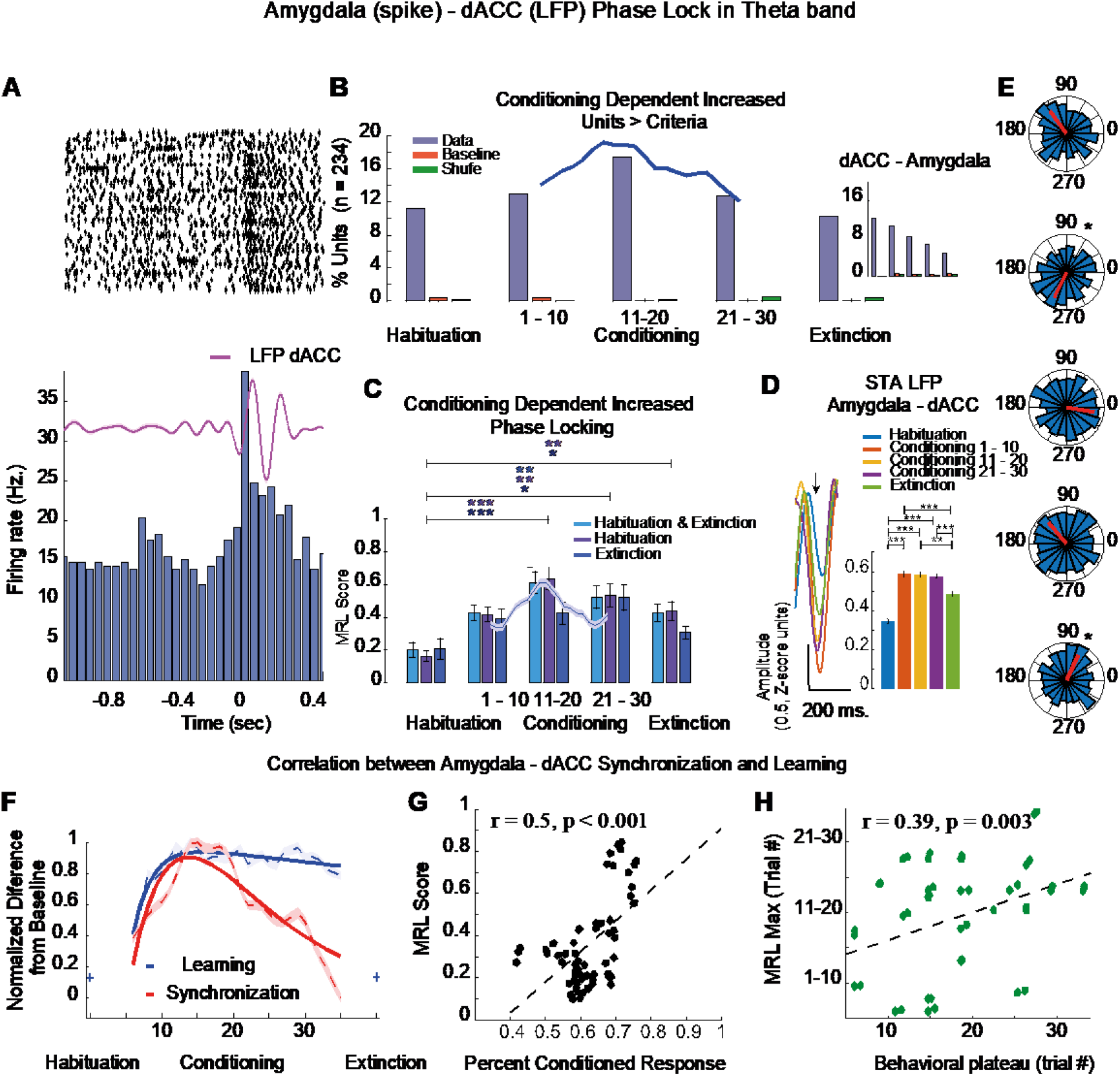
Phase locking depends on the stage of conditioning. (A) a raster plot (top) and PSTH (bottom) of amygdala spikes overlaid with mean theta activity from dACC (purple). With conditioning, dACC theta became more phase locked to amygdala spikes. (B) The proportion of phase locked amygdala units increased by 35% while that of the shuffled data and intertrial interval baseline activity did not change significantly. The solid line depicts the percent of significant amygdala using a sliding window of 5 trials. In contrast, the proportion of dACC phase-locked units decreased during conditioning by 32% (inset). (C) Synchrony score (MRL) increased during conditioning in pairs of electrodes that exhibited non-significant synchrony during habituation and extinction (n = 33), habituation only (n = 28) or extinction only (n = 32). Purple solid line shows the change in habituation only units using a sliding window of 5 trials. (D) an example of spike-triggered-averaging (STA) of theta activity averaged on spikes taken from the 500 ms. post-CS, and the averaged z-score separately for the different learning phases (one way ANOVA for learning stage, *F* = 7.84, *df* = 4, *p* = 3^−6^, post-hoc analysis results are displayed as asterisks. (E) Distribution of angle/phase of amygdala spikes (significance phase was observed in trials 1-10 of conditioning and of extinction learning). (F) Behavioral learning curve (blue dashed line) and the percent of significant units (red dashed line). Solid lines represent exponent fit (Learning: r^2^ = 0.86, RMSE = 0.04, Synchrony: r^2^ = 0.83, RMSE = 0.11). (G) Co-variation of synchrony magnitude and behavioral performance. (H) Number-of-trials to peak performance and to peak synchrony, per-session. * *p* < 0.05, ** *p* < 0.01, *** *p* < 0.001.

### Phase locking is inversely correlated with performance

The amygdala – dACC phase synchrony increased during initial stages of conditioning and started to decline during later trials, when behavior reached plateau and CS-US associations are already formed (Fig. 3f). In order to examine the co-variation of synchrony and behavioral conditioned responses per daily session, we correlated the mean synchrony score in pairs of electrodes during the first 20 trials of acquisition with the conditioned response during these trials. We found a positive relationship (Fig. 3g, Spearman’s correlation, n = 65, *r* = 0.5, *p* = 6^−5^), suggesting that inter-area synchrony contributes to the formation of the associations. To further examine if the synchrony contributes more to the acquisition of CRs or to their maintenance, we correlated the day by day number-of-trials to peak in synchrony with the number-of-trials to behavioral plateau. The positive correlation provides further evidence that the cross-regional synchrony is related to the error-signal and CR acquisition (McMains and Kastner, 2011; Rutishauser et al., 2010) (fig. 3h, Spearman’s correlation, n = 65, *r* = 0.39, *p* = 0.003, median trial to peak behavior = 11.1 ± 1.07 SE, median to peak MRL = 14 ± 0.96).

## Discussion

In the present study we show for the first time the importance of theta oscillations in the primate amygdala – dACC circuit during aversive learning. We find within-regional phase reset and further find inter-regional phase synchronization, where phase resetting in the amygdala was followed by a reset in the dACC. The oscillatory phase synchronization was accompanied by phase locking of dACC theta oscillations to amygdala spikes, but not vice versa. These findings expand on previous results in the rodent amygdala – prefrontal circuit (Adhikari et al., 2010; Berry and Seager, 2001; Courtin et al., 2014; Karalis et al., 2016; Likhtik et al., 2014; Popa et al., 2010) and on cortico-cortical communication findings in primates (Jutras and Buffalo, 2010; Liebe et al., 2012; Rutishauser et al., 2010). Together, our results point to a specific circuit and oscillatory mechanism in the primate brain that contributes to learning and formation of aversive memories.

The findings support a local and circuit level role for theta synchronization in the primate brain (Lee et al., 2005; Paz et al., 2008; Seidenbecher et al., 2003). Although theta phase resetting was shown to facilitate neuronal responses to sensory stimuli in the primate (Lakatos et al., 2007) and was suggested to optimize the processing of incoming information (Jutras and Buffalo, 2010), the functional role during learning was demonstrated only in cortico-cortical circuits and not in amygdala – prefrontal circuits. Moreover, whereas in cortico-cortical circuits synchrony was associated with memory strength; here we provide evidence that it is associated with acquisition of aversive associations and the representation of error-signal and its dynamics (Li et al., 2011). The conditioning-dependent phase resetting in the amygdala and dACC and phase locking between amygdala spikes and theta oscillations in the dACC extends the notion that memory formation relies on amygdala inputs to prefrontal regions (Burgos-Robles et al., 2017; Klavir et al., 2017; Senn et al., 2014; Sotres-Bayon et al., 2012) and on theta synchronization (Rutishauser et al., 2010, 2015), and provide first direct evidence for such a circuit level mechanism in the primate, as well as describe its dynamics during learning.

### Development of theta phase resetting in the amygdala and dACC during aversive conditioning

Theta oscillations are associated with active processing of information and stimulus-related information (Buzsáki, 2002; Jones and Wilson, 2005; Jutras et al., 2013), and several recent rodent studies suggested a functional role for oscillatory phase synchronization in spatial behavior, motor and emotional conditioning and memory consolidation (Berry and Seager, 2001; Buzsáki, 2002; Karalis et al., 2016; Likhtik et al., 2014; Popa et al., 2010). In the amygdala and medial prefrontal cortex of rodents, theta power and synchrony were shown to increase in response to tone conditioned stimulus and just ahead of the expected unconditioned aversive stimulus (Adhikari et al., 2010; Courtin et al., 2014; Paré and Collins, 2000; Paz et al., 2008; Seidenbecher et al., 2003). In line with these findings, we found increased theta power and a development of theta oscillations phase reset in the amygdala and dACC of primates during aversive conditioning. Taken together, the cross-species phase resetting suggests a functional mechanism that contributes to learning and a general phenomenon in fear learning which is robust across circuits.

### Inter-area synchrony increases during conditioning

Accumulating evidence has suggested that behavior demands coordinated activity among groups of distant neurons. In line with this suggestion, hippocampal – medial prefrontal cortex synchrony was positively correlate with anxiety behavior in mice (Adhikari et al., 2010), and increased amygdala – prefrontal theta synchrony was reported in response to the CS^+^ but not to the CS^-^ in another rodent study (Likhtik et al., 2014). Here, we observed increased theta synchrony between the amygdala and dACC of primates that emerged during fear conditioning. Similar to previous findings in the oscillation domain in rodents (Likhtik et al., 2014) and spike domain in primates (Klavir et al., 2013), we found a clear amygdala to dACC directionality in theta oscillatory activity in response to the CS. This finding is also in line with the hippocampus-to-prefrontal direction of information transfer reported previously in rodents (Jones and Wilson, 2005; Siapas et al., 2005), and suggests a similar mechanism in the amygdala-to-prefrontal circuit in primates.

The temporal lag between theta oscillations in the amygdala and that of the dACC was at the range of 10-20 ms, in both measures used in this study (the cross-correlation and *MRL*). Such a delay is likely to imply inter-area communication mechanism rather than a shared common input (Fries, 2005; Liebe et al., 2012). Taken together, the increased inter-area oscillatory phase synchronization can serve as a mean for reducing trial-by-trial variability and facilitating inter-area communication.

### Spike – LFP synchronization

It was recently suggested that the theta oscillations are used to organize and synchronize inter-area spiking activity (Benchenane et al., 2010; Courtin et al., 2014). Our results indicate that this can be the case in the primate brain as well, due to the fact that we found that amygdala spikes were more phase locked to dACC theta oscillations than vice versa. Our result is also consistent with visual (v4) - prefrontal cortex spike - LFP phase locking reported in primates (Liebe et al., 2012) and with the amygdala to dACC directionality of spiking activity we reported in the past (Klavir et al., 2013), but unlike a recent work in rodents (Likhtik et al., 2014). Such an amygdala-to-dACC mechanism can suggest that amygdala output is more effective in, and is used to drive oscillatory activity and synchronize activity in the dACC. We previously showed that in primates, spiking synchrony in the amygdala - prefrontal network underlies persistent aversive memories (Livneh and Paz, 2012). Therefore, we now suggest that presynaptic amygdala output can serve to drive postsynaptic oscillations in the dACC and increase the probability for spike in the dACC. Such a well-timed synchronized activity can facilitate learning and memory through decreased trial-by-trial variability (Darling et al., 2011; Taub et al., 2014), increased postsynaptic excitability (Sehgal et al., 2014; Yiu et al., 2014) or both.

### Synchrony as an error-signal transfer mechanism during learning

We found that synchrony between the amygdala and the dACC first increased during initial stages of conditioning, and the strong positive relationship found between synchrony and performance suggests that inter-area synchronization contributes to development of behavioral performance. This finding is in line with previous findings in non-human and human primates (Liebe et al., 2012; Rutishauser et al., 2010). Yet, whereas such mechanisms were documented only in cortico-cortical circuits (Liebe et al., 2012), our work demonstrates that this mechanism underlies similar function in a subcortical – cortical circuits, and for aversive learning. Interestingly, we found that synchrony started to decline after behavior reached a plateau, yet when paired CS-US trials are still presented. This finding points to an error-signal that is higher early in learning and diminishes as it progresses, and suggests that it is mediated at least in part by theta oscillations in the primate amygdala-prefrontal circuit (Costa et al., 2016). This finding might imply that synchrony in the amygdala – dACC circuit contributes more to acquisition of new behaviors and less for maintenance (Li et al., 2011; Roesch et al., 2010; Rudebeck et al., 2013). Finally, our findings support the assumption that learning and memory formation relies on spike – theta synchronization (Rutishauser et al., 2010, 2015).

In conclusion, we show that theta oscillations serve as a neuronal mechanism for long range communication and directional information transfer between subcortical and cortical structures during emotional learning in the primate. This mechanism can further be used to suggest a specific target for intervention in human psychopathologies (Averbeck and Chafee, 2016; Etkin et al., 2015).

## Author Contributions

A.T. and R.P. designed and performed the study. A.T. analyzed the data. A.T. and R.P. wrote the manuscript.

## Acknowledgments

We are grateful to Dr. Uri Livneh for experimental design and data collection. We thank Yossi Shohat for major contribution to the work and welfare of the animals; Dr. Eilat Kahana for help with medical and surgical procedures; Dr. Edna Furman-Haran and Nachum Stern for MRI procedures. This work was supported by ISF #26613 and ERC-FP7-StG #281171 grants to R. Paz

## STAR Methods

### Animals

Two male macaca fascicularis (4–7 kg) were implanted with a recording chamber above the right amygdala and dACC. All surgical and experimental procedures were approved and conducted in accordance with the regulations of the Weizmann Institute Animal Care and Use Committee (IACUC), following NIH regulations and with AAALAC accreditation. Anatomical magnetic resonance imaging (MRI) scans with positioning electrodes were acquired before, during and after the recording period using a 3-T MRI scanner (MAGNETOM Trio, Siemens). Scans were used to determine the depth and location of the regions.

### Recordings

Each day, three to six microelectrodes (0.6–1.2 MΩ glass/narylene-coated tungsten, Alpha Omega or We-Sense) were lowered through a metal guide (Gauge 25xxtw, OD: 0.51 mm, ID: 0.41 mm, Cadence) into the amygdala and the dACC according to electrophysiological and anatomical markers using a head tower and electrode-positioning system (Alpha Omega). Electrode signals were preamplified, 0.3–6 kHz band-pass filtered, and sampled at 25 kHz, and online spike sorting was performed using a template-based algorithm (Alpha Lab Pro, Alpha Omega). At the end of the recording period, offline spike sorting was performed for all sessions to improve unit isolation (offline sorter, Plexon). For LFP signal, signals were preamplified, 0.5 – 125 Hz band-pass filtered, and sampled at 781.25 Hz.

### Behavior

Each daily experimental session was consisted of an habituation phase of ten presentations of the pure tone CS (randomly chosen from 1,000–2,400 Hz.) that was followed by acquisition phase of 30 trials of paired CS-US, when the US is an aversive odor (propionic acid diluted in mineral oil; Sigma-Aldrich). CS was triggered by realtime detection of inhale onset, and odor (US) was released at the following inhale onset (but not before 1000 ms elapsed). Twenty unpaired CSs were used in order to extinguish the acquired CS – US association. Extinction data was analyzed for the last ten trials in order to ensure full extinction of behavioral and neuronal CRs(Livneh and Paz, 2012). Behavioral responses were computed as the area under the curve 0–350 ms. from inhae onset and normalized by the habituation inhale volume.

### Data Analysis

Data were analyzed using a combination of custom-written MATLAB scripts (MathWorks, MA) and Circular Statistics Toolbox(Berens, 2009). To evaluate LFP power across different frequencies we looked at spectrograms by obtaining the multitaper power spectral density estimation (Thomson multitaper method) of unfiltered LFP data during one second during baseline and CS periods (781 samples, time-bandwidth product of 4 and NFFT of 1024). Log-log change in LFP power was computed for CS period as above and mean of five consecutive seconds during baseline activity.

Spike autocorrelation was calculated by computing the temporal autocorrelation of the spike train during 500 and 1000 ms. following CS onset or during baseline activity. Spikes were binned (5 ms.) and number of spikes was normalization by the number of bins, with the center peak removed. Increased in theta autocorrelation was defined by calculating the power spectral density estimate of the cross-correlation vector for CS and baseline.

Estimation of change in theta power during conditioning was done for each trial using a multitaper power spectral density estimation as above, during 3 sec. ahead and 2 sec. following the CS onset, with sliding window of 50 ms. Power spectral density estimation was done separately for pre and post CS to avoid smearing of changes in theta power between pre- and post-CS time bins. Following the above, power was summed and a mean was calculated for power between 4 - 8 Hz. In order to compute the absolute change in theta power in response to the CS along conditioning, we calculated the relative change in theta power in response to the CS vs. pre-CS using the following:

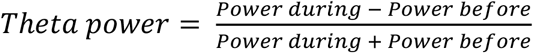

Estimation of LFP phase - phase synchrony was done by extracting phase information from the Hilbert transform of the theta-filtered (4 - 8 Hz, FIR filter) signal. We next generated histograms of the phase distributions and the half-width of these distributions was calculated from the circular standard deviation.

Phase - phase locking value was defined as the mean resultant length (MRL) and was calculated using the following

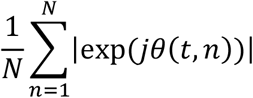

where θ(t, n) donates the phase difference ϕ1(t, n) - ϕ 2(t, n)(Lachaux et al., 1999). Thus, MRL values ranged between zero (no phase-locking) and one (perfect phase-locking).

For analyzing inter-area power correlation we computed the mean theta power for each trials as above and calculated the mean Spearman’s correlation averaged over group of ten successive trials. Differences in correlation scores were computed using Fisher r-to-z transformation.

To analyze the directionality of amygdala – dACC activity we calculated phase – phase MRL as a function of lag between the amygdala and dACC for -200 to 200 ms. centered at CS onset in steps of 10 ms.

Inter-area crosscorrelation was calculated by computing the mean of the correlation during one second starting at CS onset with maximal lag of 100 ms. for each consecutive 10 trials (i.e., habituation, three stages of conditioning and last ten trials of extinction) and using a running window of 5 trials with 4 trials overlap.

For calculation of imaginary coherence, we computed the absolute imaginary cross power spectral density estimate using Welch’s method over one second starting at CS onset using the following

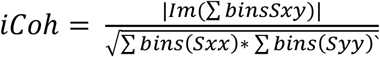

where Sxy donates the cross-spectrum, Sxx and Syy donates the auto-spectra and summation is done over the spectrogram bins corresponding to the quantified time bin(Karalis et al., 2016; Nolte et al., 2004).

For spike – LFP phase synchrony, MRL value was calculated using the above by assigning each spike event during 500 ms. after CS onset and during baseline its corresponding theta phase value. Spike – LFP phase MRL was calculated for each consecutive 10 trials and for running window of 5 trials with 4 trials overlap. MRL of shuffled data was obtained by shuffling the location of spikes in time during the CS 1000 times. Criteria for significant MRL value were obtained by calculating the MRL value for 10 randomly drawn normally distributed numbers between ±Pi for 10000 times. Criteria for significant MRL value were defined as > 95^th^ percentile of the distribution.

For percent units above criteria we computed the percent of amygdala spike – dACC LFP pair of electrodes with averaged MRL above criteria. MRL was computed over 10 consecutive trials i.e., habituation, conditioning 1 – 10, 11 – 20, 21 – 30 and extinction.

Conditioning dependent change in spike – LFP synchrony was computed for pairs of electrodes with MRL below criteria either during habituation, extinction or habituation and extinction. We included only pairs of electrodes that exhibited MRL above criteria during at list one of the three epochs of conditioning trials, i.e., conditioning 1 – 10, 11 – 20 or 21 – 30.

For synchrony – percent CR correlation we calculated the mean daily MRL score using a running window of 5 trials with overlap of 4 trials over the first 20 trials. Behavioral performance was calculated as percent trials with significant response to the CS.

For correlation between peak MRL and behavioral plateau we correlated the day by day number of trials for peak in amygdala-dACC synchronization with the number of trials for obtaining a maximal percent conditioned response.

## References

Adhikari, A., Topiwala, M.A., and Gordon, J.A. (2010). Synchronized Activity between the Ventral Hippocampus and the Medial Prefrontal Cortex during Anxiety. Neuron 65, 257–269.

Averbeck, B.B., and Chafee, M.V. (2016). Using model systems to understand errant plasticity mechanisms in psychiatric disorders. Nat. Neurosci. 19, 1418–1425.

Benchenane, K., Peyrache, A., Khamassi, M., Tierney, P.L., Gioanni, Y., Battaglia, F.P., and Wiener, S.I. (2010). Coherent Theta Oscillations and Reorganization of Spike Timing in the Hippocampal-Prefrontal Network upon Learning. Neuron 66, 921–936.

Berens P. (2009). CircStat: A MATLAB Toolbox for Circular Statistics. J. Stat. Softw. 31, 1–21.

Berry, S.D., and Seager, M.A. (2001). Hippocampal Theta Oscillations and Classical Conditioning. Neurobiol. Learn. Mem. 76, 298–313.

Burgos-Robles, A., Kimchi, E.Y., Izadmehr, E.M., Porzenheim, M.J., Ramos-Guasp, W.A., Nieh, E.H., Felix-Ortiz, A.C., Namburi, P., Leppla, C.A., Presbrey, K.N., et al. (2017). Amygdala inputs to prefrontal cortex guide behavior amid conflicting cues of reward and punishment. Nat. Neurosci. 20, 824–835.

Buzsáki G. (2002). Theta Oscillations in the Hippocampus. Neuron 33, 325–340.

Buzsáki, G., Anastassiou, C.A., and Koch, C. (2012). The origin of extracellular fields and currents — EEG, ECoG, LFP and spikes. Nat. Rev. Neurosci. 13, 407–420.

Costa, V.D., Dal Monte, O., Lucas, D.R., Murray, E.A., and Averbeck, B.B. (2016). Amygdala and Ventral Striatum Make Distinct Contributions to Reinforcement Learning. Neuron 92, 505–517.

Courtin, J., Chaudun, F., Rozeske, R.R., Karalis, N., Gonzalez-Campo, C., Wurtz, H., Abdi, A., Baufreton, J., Bienvenu, T.C.M., and Herry, C. (2014). Prefrontal parvalbumin interneurons shape neuronal activity to drive fear expression. Nature 505, 92–96.

Darling, R.D., Takatsuki, K., Griffin, A.L., and Berry, S.D. (2011). Eyeblink conditioning contingent on hippocampal theta enhances hippocampal and medial prefrontal responses. J. Neurophysiol. 105, 2213–2224.

Deffains, M., Iskhakova, L., Katabi, S., Haber, S.N., Israel, Z., and Bergman, H. (2016). Subthalamic, not striatal, activity correlates with basal ganglia downstream activity in normal and parkinsonian monkeys. eLife 5, 5.

Etkin, A., Büchel, C., and Gross, J.J. (2015). The neural bases of emotion regulation. Nat. Rev. Neurosci. 16, 693–700.

Fries P. (2005). A mechanism for cognitive dynamics: neuronal communication through neuronal coherence. Trends Cogn. Sci. 9, 474–480.

Jones, M.W., and Wilson, M.A. (2005). Theta Rhythms Coordinate Hippocampal-Prefrontal Interactions in a Spatial Memory Task. PLOS Biol. 3, e402.

Jutras, M.J., and Buffalo, E.A. (2010). Synchronous neural activity and memory formation. Curr. Opin. Neurobiol. 20, 150–155.

Jutras, M.J., Fries, P., and Buffalo, E.A. (2013). Oscillatory activity in the monkey hippocampus during visual exploration and memory formation. Proc. Natl. Acad. Sci. 110, 13144–13149.

Karalis, N., Dejean, C., Chaudun, F., Khoder, S., Rozeske, R.R., Wurtz, H., Bagur, S., Benchenane, K., Sirota, A., Courtin, J., et al. (2016). 4-Hz oscillations synchronize prefrontal-amygdala circuits during fear behavior. Nat. Neurosci. 19, 605–612.

Klavir, O., Genud-Gabai, R., and Paz, R. (2013). Functional connectivity between amygdala and cingulate cortex for adaptive aversive learning. Neuron 80, 1290–1300.

Klavir, O., Prigge, M., Sarel, A., Paz, R., and Yizhar, O. (2017). Manipulating fear associations via optogenetic modulation of amygdala inputs to prefrontal cortex. Nat. Neurosci. 20, 836844.

Lachaux, J.P., Rodriguez, E., Martinerie, J., and Varela, F.J. (1999). Measuring phase synchrony in brain signals. Hum. Brain Mapp. 8, 194–208.

Lakatos, P., Chen, C.-M., O’Connell, M.N., Mills, A., and Schroeder, C.E. (2007). Neuronal Oscillations and Multisensory Interaction in Primary Auditory Cortex. Neuron 53, 279–292.

Lee, H., Simpson, G.V., Logothetis, N.K., and Rainer, G. (2005). Phase Locking of Single Neuron Activity to Theta Oscillations during Working Memory in Monkey Extrastriate Visual Cortex. Neuron 45, 147–156.

Li, J., Schiller, D., Schoenbaum, G., Phelps, E.A., and Daw, N.D. (2011). Differential roles of human striatum and amygdala in associative learning. Nat. Neurosci. 14, 1250–1252.

Liebe, S., Hoerzer, G.M., Logothetis, N.K., and Rainer, G. (2012). Theta coupling between V4 and prefrontal cortex predicts visual short-term memory performance. Nat. Neurosci. 15, 456–462.

Likhtik, E., Stujenske, J.M., Topiwala, M.A., Harris, A.Z., and Gordon, J.A. (2014). Prefrontal entrainment of amygdala activity signals safety in learned fear and innate anxiety. Nat. Neurosci. 17, 106–113.

Livneh, U., and Paz, R. (2012). Amygdala-prefrontal synchronization underlies resistance to extinction of aversive memories. Neuron 75, 133–142.

McMains, S., and Kastner, S. (2011). Interactions of top-down and bottom-up mechanisms in human visual cortex. J. Neurosci. 31, 587–597.

Nolte, G., Bai, O., Wheaton, L., Mari, Z., Vorbach, S., and Hallett, M. (2004). Identifying true brain interaction from EEG data using the imaginary part of coherency. Clin. Neurophysiol. 115, 2292–2307.

Paré, D., and Collins, D.R. (2000). Neuronal Correlates of Fear in the Lateral Amygdala: Multiple Extracellular Recordings in Conscious Cats. J. Neurosci. 20, 2701–2710.

Paz R., Bauer E.P., and Paré, D. (2008). Theta synchronizes the activity of medial prefrontal neurons during learning. Learn. Mem. 15, 524–531.

Popa D., Duvarci S., Popescu A.T., Léna C., and Paré, D. (2010). Coherent amygdalocortical theta promotes fear memory consolidation during paradoxical sleep. Proc. Natl. Acad. Sci. U. S. A. 107, 6516–6519.

Roesch, M.R., Calu, D.J., Esber, G.R., and Schoenbaum, G. (2010). Neural Correlates of Variations in Event Processing during Learning in Basolateral Amygdala. J. Neurosci. 30, 2464–2471.

Rudebeck, P.H., Mitz, A.R., Chacko, R.V., and Murray, E.A. (2013). Effects of Amygdala Lesions on Reward-Value Coding in Orbital and Medial Prefrontal Cortex. Neuron 80, 15191531.

Rutishauser, U., Ross, I.B., Mamelak, A.N., and Schuman, E.M. (2010). Human memory strength is predicted by theta-frequency phase-locking of single neurons. Nature 464, 903–907.

Rutishauser, U., Mamelak, A.N., and Adolphs, R. (2015). The primate amygdala in social perception – insights from electrophysiological recordings and stimulation. Trends Neurosci. 38, 295–306.

Sehgal, M., Ehlers, V.L., and Moyer, J.R. (2014). Learning enhances intrinsic excitability in a subset of lateral amygdala neurons. Learn. Mem. 21, 161–170.

Seidenbecher, T., Laxmi, T.R., Stork, O., and Pape, H.-C. (2003). Amygdalar and Hippocampal Theta Rhythm Synchronization During Fear Memory Retrieval. Science 301, 846–850.

Senn, V., Wolff, S.B.E., Herry, C., Grenier, F., Ehrlich, I., Gründemann, J., Fadok, J.P., Müller, C., Letzkus, J.J., and Lüthi, A. (2014). Long-Range Connectivity Defines Behavioral Specificity of Amygdala Neurons. Neuron 81, 428–437.

Siapas, A.G., Lubenov, E.V., and Wilson, M.A. (2005). Prefrontal phase locking to hippocampal theta oscillations. Neuron 46, 141–151.

Sotres-Bayon, F., Sierra-Mercado, D., Pardilla-Delgado, E., and Quirk, G.J. (2012). Gating of Fear in Prelimbic Cortex by Hippocampal and Amygdala Inputs. Neuron 76, 804–812.

Taub, A.H., Segalis, E., Marcus-Kalish, M., and Mintz, M. (2014). Acceleration of cerebellar conditioning through improved detection of its sensory input. Brain-Comput. Interfaces 1, 516.

Yiu, A.P., Mercaldo, V., Yan, C., Richards, B., Rashid, A.J., Hsiang, H.-L.L., Pressey, J., Mahadevan, V., Tran, M.M., Kushner, S.A., et al. (2014). Neurons Are Recruited to a Memory Trace Based on Relative Neuronal Excitability Immediately before Training. Neuron 83, 722735.

